# Spatio-temporal mapping of mechanical force generated by macrophages during FcγR-dependent phagocytosis reveals adaptation to target stiffness

**DOI:** 10.1101/2020.04.14.041335

**Authors:** Pablo Rougerie, Dianne Cox

## Abstract

Macrophage phagocytosis is a strikingly flexible process central to pathogen clearance and is an attractive target for the development of anti-cancer immunotherapies. To harness the adaptability of phagocytosis, we must understand how macrophages can successfully deform their plasma membrane. While the signaling pathways and the molecular motors responsible for this deformation have been studied for many years, we only have limited insight into the mechanics that drive the formation of the phagocytic cup. Using Traction Force Microscopy (TFM), we have been able to characterize the spatio-temporal dynamics of mechanical forces generated in the course of FcγR-dependent frustrated phagocytosis and we determined whether this was affected by the stiffness of the potential phagocytic targets. We observed that frustrated phagocytosis is an atypical form of spreading where the cell deformation rate is unaffected by the substrate stiffness. Interestingly, the cell initially extends without forces being recorded then switches to a mode of pseudopod extension involving spatially organized force transmission. Importantly we demonstrate that macrophages adapt to the substrate stiffness primarily through a modulation of the magnitude of mechanical stress exerted, and not through modification of the mechanical stress kinetics or distribution. Altogether, we suggest that macrophage phagocytosis exhibits a clear resilience to variations of the phagocytic target stiffness and this is favored by an adaptation of their mechanical response.

## INTRODUCTION

Phagocytosis is a widespread contact-induced phenomenon through which cells can engulf solid particles greater than 0.5μm in size. The phagocytic activity of leukocytes, such as macrophages, is crucial for pathogen clearance (Donnelly & Barnes 2012). Scavenging macrophages are also responsible for the removal of necrotic and apoptotic cells, preventing the onset of inflammation and autoimmune disease (Elliott & Ravichandran 2010). Interestingly, multiple examples also show that macrophage-mediated phagocytosis can be harnessed for immunotherapy in order to foster the elimination of opsonized, living cancer cells (Gul et al 2014, Weiskopf et al 2013). Macrophages express a variety of phagocytic receptors including the well-characterized Fcγ receptor (FcγR) that binds to the Fc portion of IgG. Upon binding of a phagocytic ligand to a cellular phagocytic receptor, the macrophage forms a membrane protrusion called the phagocytic cup. The phagocytic cup then extends around the target and the rim of the cup eventually fuses at the distal pole, sealing the target inside (Rougerie et al 2013). Owing to the omnipresence and biomedical potential of phagocytosis, significant effort has been dedicated to understand how the phagocytic cup can successfully form, extend, and close. The phagocytic cup can morphologically be interpreted as the spreading of a cell around a three-dimensional substrate (the target). Logically, phagocytic cup formation shares signaling pathways with the more classic spreading of cells like fibroblasts on adhesive substrates. Downstream of the FcγR, signaling involving the Rho GTPases Rac or Cdc42 and phosphoinositides (Cox et al 1999, Hoppe & Swanson 2004, Swanson 2008) regulate the spatio-temporal activity of actin and myosins and induce the extension of the phagocytic cup (Freeman & Grinstein 2014, Rougerie et al 2013). Many studies have shown that cortical tension and mechanical traction forces play a central role during cell spreading (Dubin-Thaler et al 2008, Gauthier et al 2011, Nisenholz et al 2014, Pietuch & Janshoff 2013, Reinhart-King et al 2005). Similarly, physical parameters must regulate the formation of the phagocytic cup (Danuser et al 2013, Fardin et al 2010). Understanding the mechanical basis of phagocytosis is thus a complementary approach to the more classic biochemical studies and therefore can bring novel and valuable information. In particular, it would provide critical insight on the striking flexibility of phagocytosis. Indeed, phagocytic targets come with different shapes, sizes and stiffness. For instance, bacteria can be as small as 1-2um and very stiff, with Young’s modulus within the megapascal (MPa) range (Tuson et al 2012). On the other hand, cancer cells are much bigger and softer, with a Young’s modulus around one kilopascal (kPa) (Cross et al 2007, Xu et al 2012). Yet it is unclear how phagocytic cells can adapt to this diversity. We thus hypothesized that macrophages would react to targets of different stiffness by adapting the mechanical dynamics during the formation of the phagocytic cup.

Mechanical studies of phagocytosis so far have mainly focused on cortical tension. Herant et al. used micropipette aspiration to show that the cortical tension increases during neutrophil phagocytosis and mechanically limits the size of the phagocytic cup (Herant et al 2005, Herant et al 2006, Herant et al 2011). Cortical tension during macrophage phagocytosis has been linked to regulation of internal membrane trafficking and velocity of cell deformation (Masters et al 2013). While cortical tension data is readily available, results on the cell-generated mechanical forces actuating the formation of a phagocytic cup are scant. Herant et al. infer from their micropipette experiments a global estimation of a ~ 50 nN protrusion force − comparable to that exerted by neutrophils during chemokinesis (Herant et al 2005, Herant et al 2006, Smith et al 2007), yet without direct force measurement. A method for directly measuring traction forces, Traction Force Microscopy (TFM), was used on a macrophage cell line and identified a low level of traction forces throughout phagocytosis (Kovari et al 2016). Once the cell reached its maximal deformation, the authors reported a transient peak of force linked to a marked cell contraction. However, the mechano-morphological behavior of macrophages during phagocytosis remains unclear. Indeed, other studies performed with a similar experimental approach did not identify a phase of cell retraction (Beningo & Wang 2002, Cannon & Swanson 1992, Masters et al 2013). In addition, the spatiotemporal stress dynamics during the extension of the phagocytic cup needs to be more thoroughly analyzed. Finally, whether and how physical parameters of the phagocytic targets such as its stiffness affects the cell mechanical forces is still unknown. To address these issues, we set out to analyze the spatiotemporal dynamics of cell-generated mechanical forces when primary Bone Marrow derived Macrophages (BMMs) attempt to phagocytose targets of different stiffness.

## MATERIAL AND METHODS

### Buffers

Buffer with Divalent Cations (BWD) was prepared as follows: 20mM Hepes, 125mM NaCl, 5mMKCl, 5mM Glucose, 10mM NaHCO_3_, 1mMKH_2_PO_4_, 1mM CaCl_2_, 1mM MgCl_2_, pH 7.4.

### Cell culture and BMM preparation

Bone marrow was isolated from the femurs and tibia of mice and grown at 37°C with 5%CO_2_ in bacterial plastic dishes in alphaMEM with 15% FBS (Sigma), 100U/mL penicillin, 100ug/mL streptomycin (Fischer) with 360 ng/mL recombinant human CSF-1 (Chiron, Emeryville, CA) to generate BMMs. Mature BMMs were detached by incubation in PBS with 10 mM EDTA, washed and resuspended in BWD.

### Coverslip activation

Coverslips were pre-activated prior to polyacrylamide gel casting by first washing overnight in 0.2M HCl followed by rinsing with distilled water. Remaining acid was then neutralized with 0.1M NaOH for 30min and rinsed with distilled water. Coverslips were then incubated on an orbital shaker in 3-aminopropyl trimethoxysilane 0.5% for 30min and rinsed extensively with water. Then the coverslips were incubated with glutaraldehyde 0.5% for 1 hour and rinsed with distilled water and air-dried overnight.

### Gel casting

Acrylamide solution was prepared containing distilled water, Hepes pH 8 35mM (final concentration), n’-tetramethylethylene di-amine (TEMED, Sigma), acrylamide (BIORAD, 40% w/v stock solution), *n,n’*-methylene-bis-acrylamide (BIORAD, 2% w/v stock solution), N-6-((acryloyl)amino)hexanoic acid crosslinker (N6, 0.56% w/v diluted in ethanol 200 proof). The N6 crosslinker (generously provided by Daniel Hammer (University of Pennsylvania) (Hind et al 2015) contains an *n*-succinimidyl ester that can be displaced by a primary amine thereby covalently binding an amine-containing ligand to the polyacylamide gel. The percentages of acrylamide and bis-acrylamide were adjusted to modify the stiffness of the gel (Hind et al 2015, Reinhart-King et al 2003) and 500nm FluoSpheres (Life Technologies) were added as position markers. The solution was degassed for 30min at room temperature in a vacuum chamber. Polymerization was initiated with Ammonium Persulfate (0.06% final, Fisher) and a drop of acrylamide solution was then added onto a siliconized glass coverslip. Immediately, an activated coverslip was placed on top of the drop to form a flat film of uniform thickness. The coverslips and gel sandwich was then placed into a humid Argon-filled chamber and incubated for 1h at room temperature to allow for acrylamide polymerization. The siliconized coverslip was then carefully removed with a sharp razorblade, to avoid any tearing of the thin gel. The gel was then immediately rinsed with distilled water and incubated overnight at 4°C with a 1mg/mL solution of human whole IgG (CellSciences) or human Fab fragment (Rockland) in 50mM HEPES pH 8. Excess antibodies were rinsed away with distilled water and unbound N6 crosslinkers were blocked by a 30min incubation in a 1% (v/v) solution of Ethanolamine (Sigma) in 50mM HEPES pH 8 and then rinsed with distilled water, The Fab- or IgG-coated, flat, elastic acrylamide gels were stored at 4°C in PBS for up to 4 days.

### Frustrated phagocytosis

For frustrated phagocytosis on elastic gels, the surfaces were prepared as explained above. For frustrated phagocytosis on glass, the glass coverslips were coated with a 1mg/mL solution of human whole IgG or Fab fragment in PBS for 1h at room temperature and excess antibodies were removed by rinsing three times with PBS and the coverslips were then stored in PBS and used the same day. Live cell imaging was performed by placing the gel or coverslip on a heated stage of a Olympus IX71 microscope equipped with a SENSICAM cooled CCD camera. BMMs suspended in BWD were then added onto the coated substrates, and images were taken every minute with a Olympus LCPlanFI 20X objective for 30min. For fixed synchronized frustrated phagocytosis assays, the substrates (gels or coverslips) were kept on ice for 5min and suspended BMMs in cold BWD were added and incubated on ice for a further 15min. The cells that did not attach to phagocytic ligands were washed away by rinsing with ice cold BWD. The substrates were then swiftly transferred to a 37°C water bath and incubated for the indicated time before fixation with a 3.7% formaldehyde solution in BWD.

### TFM

The gels (Fab- or IgG-coated) were covered by a drop of BWD and incubated in the C0_2_-buffered, thermostated microscopic chamber. The mechanical field has been quantified as in Hind *el al.(Hind et al 2015)*. Briefly, phase images were collected every 20s or 30s under a 40X objective, for 30min. For each phase image, the corresponding fluorescent image of the gel-embedded beads field was also recorded. At the end of the acquisition, the imaging buffer was carefully replaced incubation with 0.5% SDS in BWD for 5 min to remove all of the attached cells and to relax the gel to its initial conformation and an image of the unstressed beads field was then taken. Cells that were away enough from the field edges or from neighboring cells were selected for quantification. The displacement field of the beads was measured by comparison to the image of the unstressed bead field using the custom written software LIBTRC 2.4 (generously provided by Micah Dembo, Boston University) (Dembo & Wang 1999). At one given time point, the cell image undergoes a tessellation and the average traction force exerted onto the substrate is provided by LIBTRC by resolving the double integral: 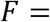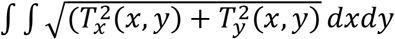 where *T_x_* (respectively *T_y_*) i the *x* (respectively *y*) component of the maximum of likelihood traction field *T* at the point of *(x,y)* coordinates. Finally, the normalization of the average force *F* by the cell area leads to the average mechanical stress given in Pa: *S=F/A* where *A* is the area of the cell.

To allow the generation of curves of the mechanical stress as a function of time, key physical values and statistical indicators of all the time frames from a single cell recording were automatically compiled into a single file using a custom written Scilab script (Scilab 5.4, Scilab Enterprises). Spatial heat maps of the local mechanical stress were generated using a custom written ImageJ (NIH) macro. In those maps, each pixel value corresponds to the mechanical stress at that same point. To further analyze the spatial organization of the force field, the areas corresponding to the top 20% pixels in each time frame of each cell were automatically selected using a custom written ImageJ (NIH) macro. For each time frame of each cell, the number of obtained “stress zones” was retrieved, and their average size and stress were calculated. Given the cell tendency to be circular during frustrated phagocytosis, we calculated the stress zones radial position by using the radial indicator 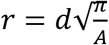, where *A* is the area of a cell at a given time point and *d* is the average distance between its stress zone centroids at that time and the cell centroid.

### Phosphotyrosine (pY) and actin staining and measurement

Fixed, synchronized phagocytosis assays were performed as mentioned above. The cells were permeabilized by a 0.2% Triton-X100 (Fisher) solution in PBS for 4min and blocked with 1% BSA (Fisher) in PBS for 30min and stained with the anti-pY primary antibody (4G10 Platinum, EMD Millipore, 1:1000) in PBS 0.2% BSA for 1h. The cells were then stained with 5ug/mL Alexa488 Donkey-anti-Mouse secondary (Jackson Immunoresearch) for 30min. F-actin was stained with 5ug/mL Alexa568-phalloidin (Molecular Probes, ThermoFischer Scientific). The cells were rinsed in PBS and imaged with a Olympus LCPlanFI 20X objective mounted on a Olympus IX71 microscope equiped with a SENSICAM cooled CCD camera. The amount of pY or F-actin was assessed by quantifying the integrated density of the fluorescence of the whole cells, as imaged at 20X.

### Statistics

One-way ANOVA and post-ANOVA Tukey tests, linear and non-linear regressions, correlation tests were performed using Prism version 5.0c.

## RESULTS

### Macrophage frustrated phagocytosis is resilient to variations in substrate stiffness

In order to investigate the effect of particle stiffness on phagocytosis, we monitored macrophage spreading on IgG-coated polyacrylamide (PA) sheets of various Young’s Moduli, a proxy for phagocytic cup formation compatible with TFM and similar to Beningo *et al* (Beningo & Wang 2002). In this so-called “frustrated phagocytosis” assay, the basal membrane of the cell shares common signaling or organization features with the phagocytic cup (Masters et al 2013) (fig 1A). We first determined whether the stiffness of the antibody-coated substrate could affect the efficiency FcγR-dependent spreading by BMMs. The range of Young’s moduli was chosen to encompass those reported in the literature for several important phagocytic targets such as cancer cells (0.5-2kPa) (Cross et al 2007, Xu et al 2012) or red blood cells (4-26kPa) (Bremmell et al 2006, Maciaszek et al 2011). Elastic gels with Young’s moduli of 1.25kPa, 2.5kPa, 7.9kPa and 15.6kPa were used and compared to glass as a reference. Upon contact with the substrate, the cells flatten and increase their contact area. While some cells merely form transient lateral protrusions without further extension, the large majority successfully enter into frustrated phagocytosis on all the stiffness conditions tested (79% to 95%), without statistically significant differences (fig 1B). Cells increased their contact area by forming a characteristic continuous, thin, adherent outer edge on all stiffness conditions (fig 1C). Cells performing frustrated phagocytosis initially displayed a phase of rapid extension before entering a second phase of lower to null velocity, regardless of substrate stiffness (fig 1D and 1E). In agreement with other studies (Masters et al 2013), the average spreading curves show that the first step is almost linear with cells rapidly increasing 4-fold within a few minutes. Run tests of linear regression shows a statistically significant departure from linearity at time = 7min (fig 1E). The second step of frustrated phagocytosis is non-linear that slows to a plateau at approximate 8-fold contact area (fig 1E). Previous observations of phagocytic and non-phagocytic spreading over glass reported a biphasic spreading kinetics with an initial fast linear spreading (termed P1) and a slowing, non-linear phase (P2) (Dubin-Thaler et al 2008, Masters et al 2013, Reinhart-King et al 2005). Our kinetics using IgG-coating acrylamide gels of different stiffness are thus analogous to the P1/P2 dynamics previously described. We did not observe sustained negative velocity at the later time points, demonstrating the absence of cell retraction (fig 1D). Importantly, BMMs were unable to undergo frustrated phagocytosis when the PA gels were coated with Fab fragment lacking the FcγR-binding Fc portion instead of full IgG (fig 1E). This indicated that the absence of the FcγR ligand prevented any spreading on the substrate. In spite of the overall consistency with previously reported spreading dynamics, macrophage frustrated phagocytosis showed one striking particularity where the kinetics of spreading was clearly conserved over all stiffnesses and the average initial velocity during the linear spreading phase was similar (fig 1D). The time at which spreading switched from the first, rapid and linear phase, to the second slow, non-linear phase was also comparable (fig 1E) as was the maximal cell area and the delay between cell landing onto the substrate and initiation of spreading (Fig 2). By contrast, numerous studies from the literature reported a positive correlation between cell spreading and the stiffness of fibronectin-, collagen- or laminin-coated substrates (Engler et al 2004, Han et al 2012, Li et al 2014, Nisenholz et al 2014, Reimer et al 2017, Tee et al 2011, Vishavkarma et al 2014, Yeung et al 2005). Therefore, the atypical resilience of FcγR-triggered spreading to substrate stiffness variation seemed a unique feature. To explain this effect, we hypothesized that macrophages were able to adapt to different stiffness by modifying the cell mechanical forces driving membrane extension during frustrated phagocytosis.

**Figure 1:**
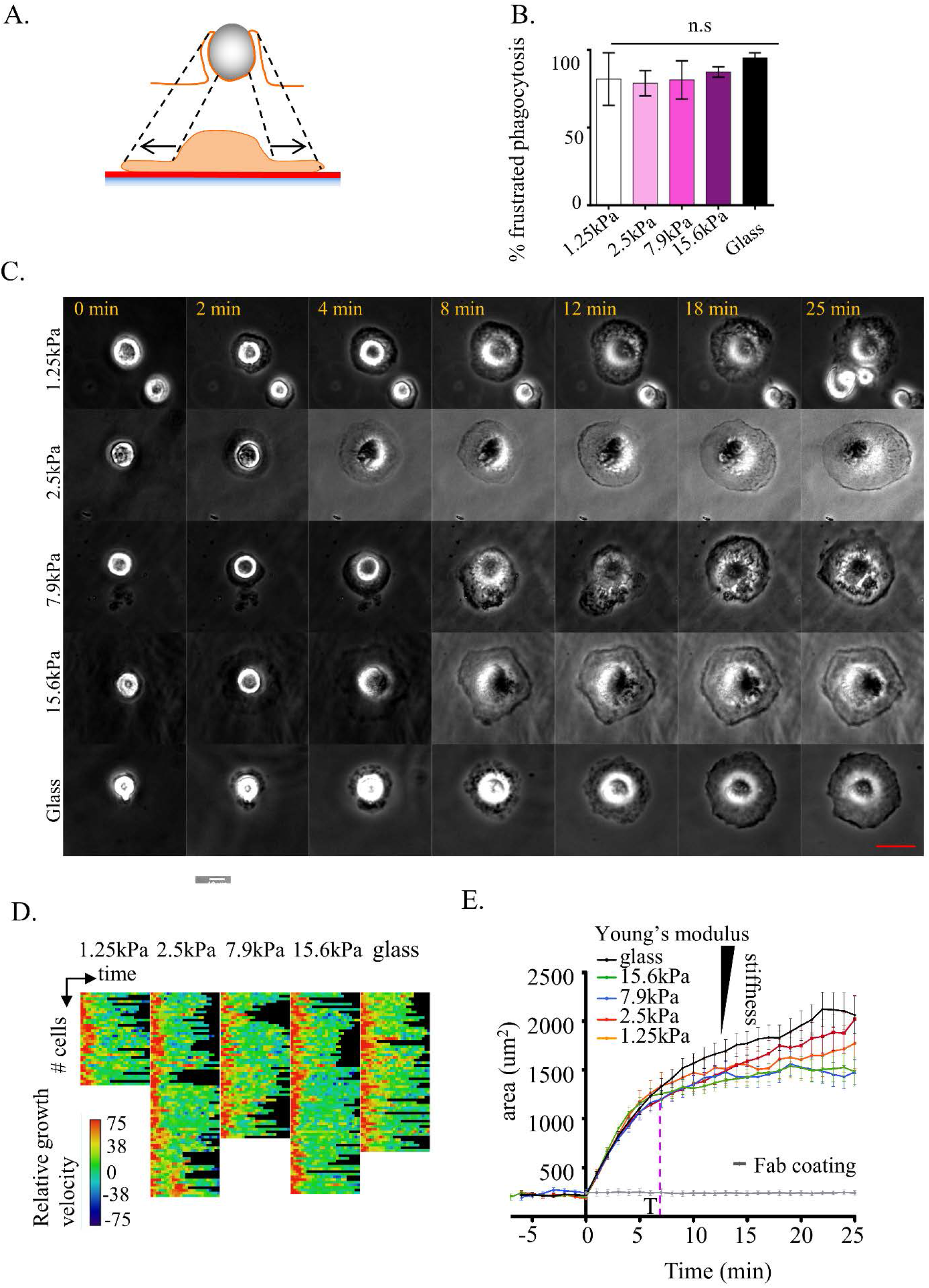
Frustrated phagocytosis is a biphasic cell spreading resilient to variations in substrate stiffness. Frustrated phagocytosis was performed on IgG-coated glass or IgG-coated PA gels of various stiffness. (A) Diagram comparing frustrated phagocytosis to traditional phagocytosis of a 3D object. Upon contact with the IgG-opsonized substrate, the cell isotropically spreads. The edges and the center of the basal membranes share molecular and mechanical analogies with the rims and the base of the classic 3D phagocytic cup, respectively. (B) Percentage of cells fully initiating frustrated phagocytosis on the indicated substrates. Cell reaching a 600um^2^ threshold (≈fold spreading) are considered having fully entered into frustrated phagocytosis, as it is associated with formation of a characteristic continuous peripheral flat protrusion (n=3 to 4 experiments. Error bar: SEM). (C) Time lapse images of macrophages during frustrated phagocytosis. One representative cell per stiffness condition. The elapsed time from the initiation of spreading is indicated. (D) Spreading cell velocity of individual cell frustrated phagocytosis. Each row represents a different cell, each column a successive time point (1min per pixel). To ease comparison, the velocities are normalized to 100% = max speed of each cell. (E) Increase of cell-substrate contact area in the course of frustrated phagocytosis. The contact areas of each cell were averaged together at each time point. The dotted magenta line indicates the spreading transition from rapid and linear to non-linearly slowing and plateauing, as determined by run test (see text). Only the time points where 5 or more cell measurements were available for a given stiffness were conserved. Only the cells showing a clear initiation of frustrated phagocytosis (cell area reaching at least 600um^2^) were considered (n=36 to 81 cells obtained from 3 to 5 independent experiments for the IgG-coated substrates, n=6 cells for the Fab negative control. Error bar: SEM).

**Figure 2:**
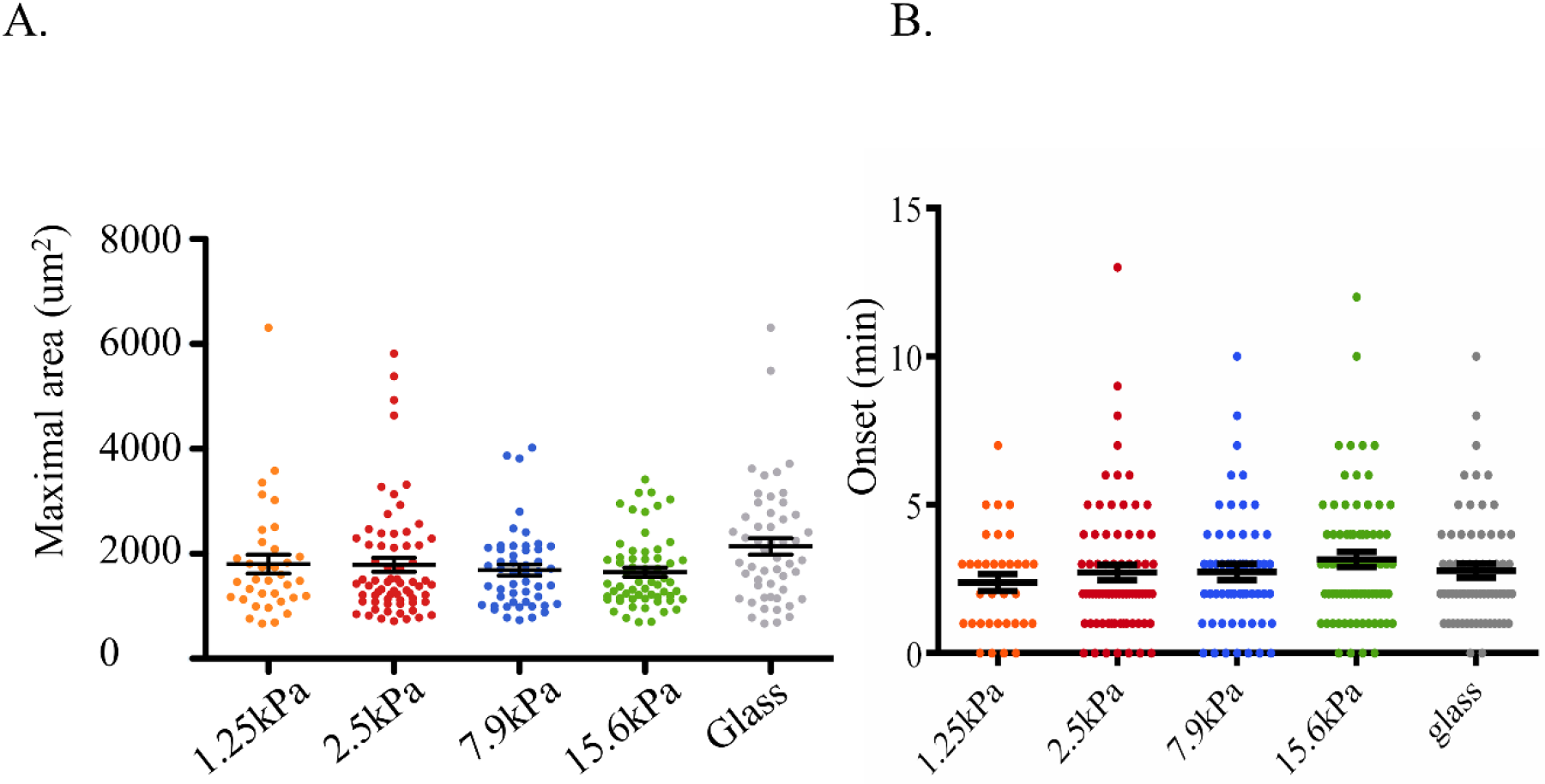
Maximal area and lag before spreading initiation show invariance to substrate stiffness. (A) Maximal area: maximal area reached by a cell at any time point of its spreading. Cells that do not reach a plateau at the end of the movie (25min) are ignored (n=34 to 68 individual cells per stiffness, obtained from 3 to 5 independent experiments. Error bar: SEM). (B) Onset: elapsed time in minutes between the moment where a given cells land on the IgG-coated substrate and the moment where it starts spreading. (n=34 to 68 individual cells per stiffness, obtained from 3 to 5 independent experiments. Error bar: SEM).

### Macrophages are mechanoresponsive to variation in phagocytic target stiffness

To investigate the potential impact of target stiffness on phagocytosis-related mechanical forces we combined the frustrated phagocytosis assay with Traction Force Microscopy (TFM) (fig 3A). TFM provides cellular mechanical force measurements with excellent resolution in space and time and has been successfully used to quantify the force field generated by cells, including primary macrophages, during migration over purely elastic PA gels (Hind et al 2015, Jannat et al 2011, Munevar et al 2001, Reinhart-King et al 2005, Reinhart-King et al 2008). When performed on IgG-coated gels, spreading caused visible localized gel deformation (fig 3A) allowing for deduction of the forces using TFM. We found that the average magnitude of mechanical stress generated by the BMMs throughout frustrated phagocytosis significantly increased with the substrate Young’s modulus (6.8Pa on 1.25kPa gel, 31.6Pa on 2.5kPa gel, 70.7Pa on 7.9kPa gel, 204.8Pa on 15.6kPa gel) (fig3B). Table 1 provides numerical values for the stress and the corresponding forces before and after the spreading transition described in figure 1. The forces were all in the nano-Newton range, reaching as high 1964nN (on a 15.6kPa gel corresponding to a 1736Pa stress). Macrophages thus showed a striking 30-fold increase between the average stress magnitude on the softest and the stiffest substrate. The correlation between cell stress and the stiffness of FcγR-binding substrate strongly fits a linear function (fig 3C). Within the range of tested values, the average mechanical stress that a BMM generated during frustrated phagocytosis was approximately 0.013 times the substrate stiffness. These results strongly demonstrated that FcγR-triggered membrane extension is not only mechanosensitive but also that the nature of its response was itself mechanical. We thus proposed that BMMs reach comparable efficiency and kinetics of frustrated phagocytosis on substrates of different stiffness through a proportional modulation of the magnitude of mechanical stress that they generate. As a consequence, the ratio between the maximal area reached by the cells and the average magnitude of their stress negatively correlate with the Young’s moduli of the gel (94.7um^2^/Pa on a 1.25kPa gel and 3.9um^2^/Pa on a 15.6kPa gel, fig 3D). Simply stated, the gain of area for every additional Pascal of stress generated was much lower at the substrate gets stiffer. Altogether, these results indicated that it is mechanically more challenging to efficiently perform frustrated phagocytosis on a stiff surface and that macrophages are able to overcome this difficulty by generating greater forces during phagocytic cup formation of stiff targets.

**Table 1:** Numeric values of the force and stress transmitted to the substrate during BMMs frustrated phagocytosis. The table indicates the minimum, average, and maximum stress and force recorded for cells either before or after the spreading transition from rapid linear spreading to slower non-linear

| Substrate<br>Young's<br>modulus<br>(kPa) | Stress (Pa) |  |  |  |  |  | Force (nN) |  |  |  |  |  |
| --- | --- | --- | --- | --- | --- | --- | --- | --- | --- | --- | --- | --- |
|  | **before** linear to non<br>linear spreading<br>transition |  |  | **after** linear to non<br>linear spreading<br>transition |  |  | **before** linear to non<br>linear spreading<br>transition |  |  | **after** linear to non<br>linear spreading<br>transition |  |  |
|  | min | averag<br>e | max | min | averag<br>e | max | min | averag<br>e | max | min | averag<br>e | max |
| 1.25 | 0 | 3.298 | 25.69 | 0 | 7.612 | 38.70 | 0 | 3.302 | 13.85 | 0 | 17.36 | 69.29 |
| 2.5 | 0 | 5.759 | 28.17 | 0 | 34.39 | 196.5 | 0 | 8.176 | 39.92 | 0 | 67.87 | 552.9 |
| 7.9 | 0 | 30.54 | 361.1 | 0 | 71.81 | 422.0 | 0 | 40.21 | 372.8 | 0 | 171.5 | 909.8 |
| 15.6 | 0 | 69.93 | 694.5 | 0 | 231.7 | 1736 | 0 | 66.35 | 738.2 | 0 | 277.7 | 1964 |

**Figure 3:**
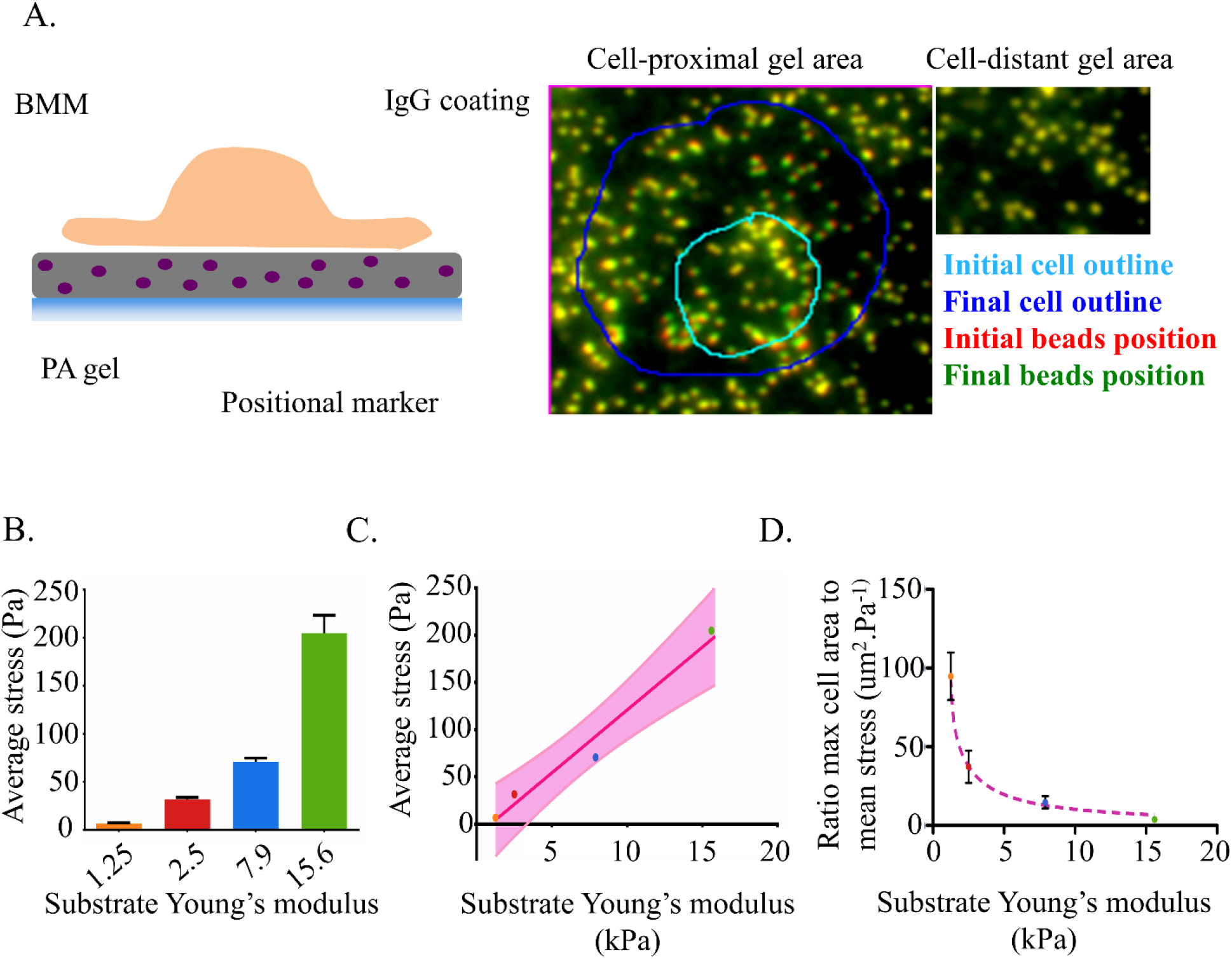
BMMs mechanically adapt to their target stiffness during frustrated phagocytosis. (A) Left: Cell Traction Forces transmitted to the substrate are detected by adding position markers to the elastic polyacrylamide gel and monitoring their displacement. Right: the cell-induced deformation of the elastic substrate is illustrated by a shift in bead positions underneath the cell at later time points. By contrast, regions away from the cells are unaffected. (B) The histogram shows the mean stress generated by cells over the entire surface from the onset of spreading to the end of recording. (1.25kPa: N = 3 cells, 2.5kPa: N =7 cells, 7.9kPa: N =9 cells, 15.6kPa: N = 7 cells, error bar: SEM). (C) The average stress delivered by cells during frustrated phagocytosis linearly correlates with the target stiffness. Regression of the stress value shown in (C). The line equation is: *y = 13.31*x − 12.17*, R^2^ = 0.9701. The 90% confidence interval is indicated. (D) The frustrated phagocytosis on stiffer substrate is mechanically more challenging as illustrated by this ratio of maximal area reached by the cell to its average mechanical stress. It fits a power series model y = 279.4x^−11.22^ + 88.55*x^−0.9350^, R^2^ = 0.7337

### Kinetics of mechanical stress generation in the course of frustrated phagocytosis

To verify that the modulation of average stress magnitude was the main mechanical adaptation to target stiffness, we examined the kinetics of stress generation during frustrated phagocytosis. Figure 4A shows the cell-substrate contact area as well as the cell-generated mechanical stress as a function of time for four representative cells (one per gel stiffness condition). The corresponding movies of these cells are shown in supplemental data. As a negative control, we looked at cells on Fab-coated 7.9kPa substrates that are unable to bind FcγR due to the lack of Fc-fragment. No mechanical stress was recorded on the Fab-coated substrate (fig 4A, red triangles) demonstrating that FcγR binding was necessary for force generation or transmission to the substrate. In the first few minutes, the cell starts to spread on IgG-coated substrates even though no stress is being recorded (fig. 4A and 4B). At the time of first mechanical stress recording, the cells have already tripled their contact area (fig 4C). This may be explained by the fact that the mechanical stress at early time points was below the detection threshold. Alternatively, it suggests the interesting possibility that the cells start extending without force transmission to the substrate before switching to a more classic extension mode based on anchoring and force transmission. Once stress began to be transmitted to the substrate, it sometimes showed significant oscillations of magnitude; in some cases, the cell returned to an unstressed situation. Eventually, the traction stress stabilized around a non-null value. Importantly, the stress magnitude reached a stable value *before* the cell was fully spread (fig 4A). The extension of the cell was thus not accompanied by a simultaneous and correlated increase of mechanical stress. Importantly, we did not find any influence of the substrate stiffness on these kinetic features of stress generation. Indeed, the delay before the first stress recording (fig 4B), the strain at the moment of first stress recording (fig 4C) and the moment of stress stabilization (Fig 4D) were all uncorrelated with substrate stiffness, with important cell to cell variation. In addition, we observed no correlation between the stress magnitude at a given time point and the corresponding spreading velocity (data not shown). The strain energy density, measuring the amount of mechanical work performed by the cell onto the substrate closely followed the evolution of the stress magnitude (fig 4E), suggesting that the cell mechanical output was mainly controlled by the stress magnitude. Therefore, it appears that the primary mechanism of mechanical adaptation to various substrate rigidities was the modulation of the overall stress magnitude showed in figure 3.

**Figure 4:**
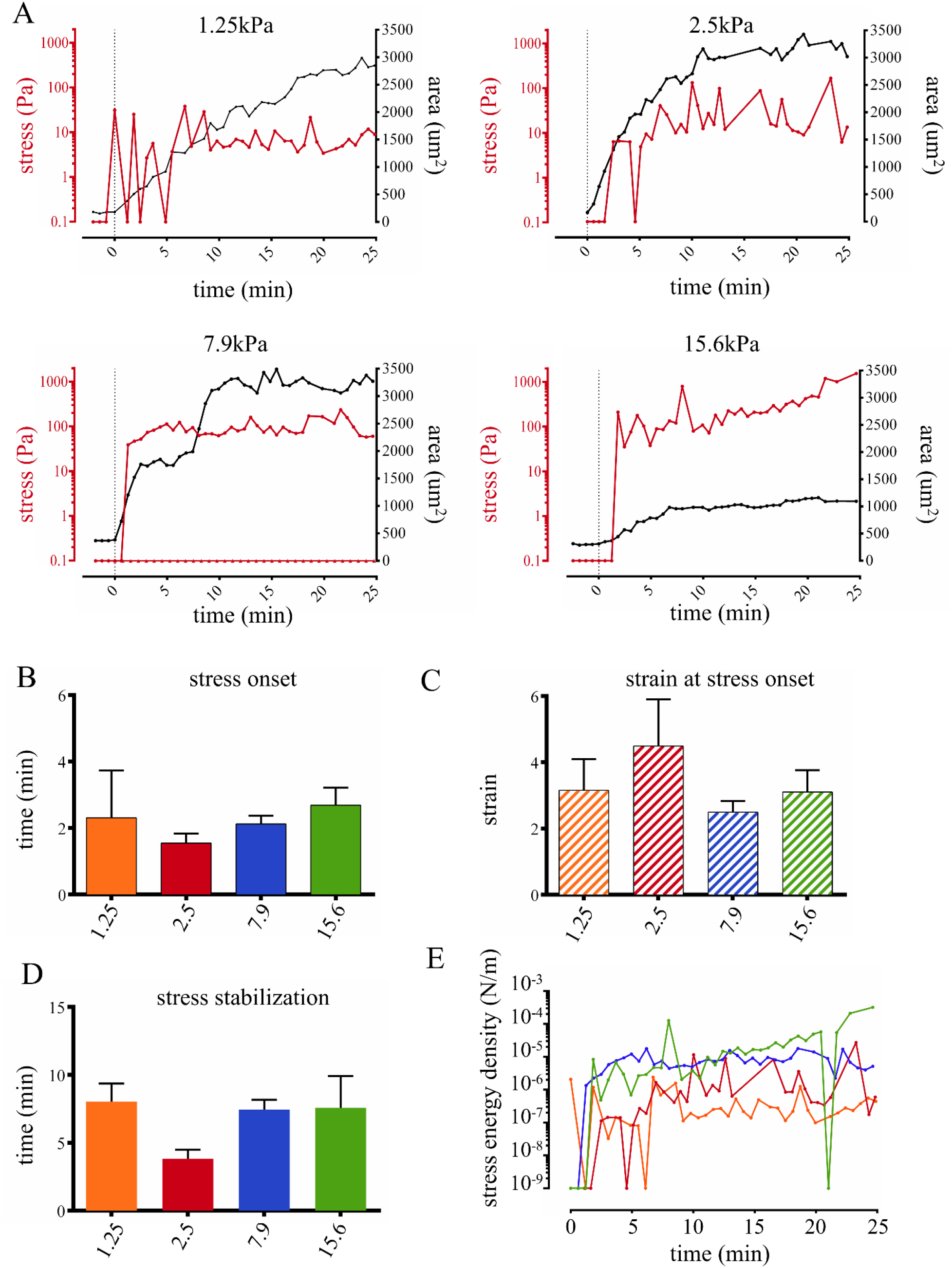
The kinetics of traction force generation is conserved over time between substrate stiffness conditions. (A) Representative recording of mechanical stress (red curve) and contact area (black curve) for one representative cell per stiffness conditions. Stress is plotted on the log left axis, the cell −substrate contact area is plotted on the right axis. The cell rapidly and linearly spreads out before transition to a slowing non-linear spreading. No traction stress is initially recorded even though the cell starts spreading. Afterward, the mechanical stress builds up and stabilizes around an average value before the cell reaches its spreading plateau. On the graph corresponding to the 7.9kPa gel condition, the triangle line show the stress measurement for cells loaded on a control Fab-coated gel (average of 8 cells). (B) Average elapsed time before the first recording of traction stress. No statistically significant difference between stiffness condition has been detected. Error bar: SEM. (C) Average strain reached by the cells at the time of first traction stress recording, highlighting that cells are able to start spreading even though the mechanical stress is either below the sensitivity threshold or not transmitted to the substrate. (D) Average time of stabilization of the mechanical stress magnitude. Error bar: SEM. (E) Corresponding strain energy density of the four cells represented in figure 4A. The strain energy density illustrate the amount of mechanical work put into the substrate by the cell traction, per unit area. Note that the strain energy density follows closely the recorded mechanical stress. The line color corresponds to the different stress conditions shown in B, C and D where orange = 1.25, red = 2.5, blue = 7.9 and green = 15.6 kPA as in the previous figures.

### Mechanical stress in the phagocytic cup is spatially organized

In many contexts such as cell spreading or migration, the cellular mechanical stress field exhibits specific spatial patterns that dictate the cell deformation and displacement (Hind et al 2015, Jannat et al 2011, Ji et al 2008, Reinhart-King et al 2005, Tanimoto & Sano 2014). Therefore, we decided to verify whether the mechanical adaptation to different stiffness was associated with modification of the spatial distribution of mechanical stress. Figure 5 shows the stress fields generated by the four representative cells used in figure 4. The corresponding time points span the entire course of frustrated phagocytosis. Additional examples (one representative cell per stiffness condition) are available in Fig S1. The stress fields over the full time course of frustrated phagocytosis are shown in movies 1 to 8. The detected stress fields were non-uniform with distinct areas of high and low stress. We termed “stress zones” as delimited subcellular areas of high stress generation in each stress field (one field per cell and per time point). Most of the time, the stress zones accumulated at the periphery of the cell and were accompanied by a large area in the center of the cell which was devoid of stress (fig 5). The corresponding stress vectors of the selected cells are represented in figure 6 indicating a clear centripetal direction of the mechanical stress (Fig 6 and Supplementary Figure S2). We then expressed the position of those stress zones relative to the cell centroid as a radial index *r*, where *r* tends toward 1 when stress zones accumulate at the periphery and toward 0 when they are positioned near the cell centroid (see *Material and Methods*). We found *r* ≈ 0.8 for all substrate stiffness confirming the peripheral restriction of stress generation (fig 7A) independently of the substrate Young’s modulus. The level of fragmentation of the stress field, as illustrated by the number of distinct stress zones in each map was comparable between the different gel stiffnesses (fig 7B). However, we detected that the level of fragmentation of the stress map was associated with an overall higher mechanical stress being generated and that this effect was more pronounced as the substrate got stiffer (fig 7C). Finally, we confirmed the tendency of mechanical stress to point inward by calculating the cosine of the stress vector at one point of the cell area with the segment formed by this point and the cell centroid (fig 7D). Thus, an average cosine of 1 would represent perfectly centripetal stress at all points and a cosine of −1 perfectly centrifuge stress. The average cosine (averaged over all time points of all cells) is above 0, confirming the dominant centripetal orientation of mechanical stress without significant effect from the substrate stiffness. Considering together the segmentation of the stress field into distinct stress zones, their accumulation at the periphery of the cell, and the clear centripetal tractions exerted at the cell edge, we concluded that the basal membrane during frustrated phagocytosis is a mechanically organized structure where cells generated a ring of traction forces at the cell edge. Although we observed that the substrate stiffness does have a slight influence on how the fragmentation of stress zones relates to higher mechanical stress, it appears that the main spatial characteristics of stress generation during frustrated phagocytosis (number and localization of stress zones, orientation of the forces) are conserved between the gel stiffness conditions. We thus drew our general conclusion that the resilience of frustrated phagocytosis to variation of target stiffness observed in figure 1 could be explained by a mechanoresponse based primarily on a modulation of the magnitude of stress being generated (fig3) whereas the time kinetics (fig 4) and spatial distribution (fig 5, 6, 7) were preserved.

**Figure 5:**
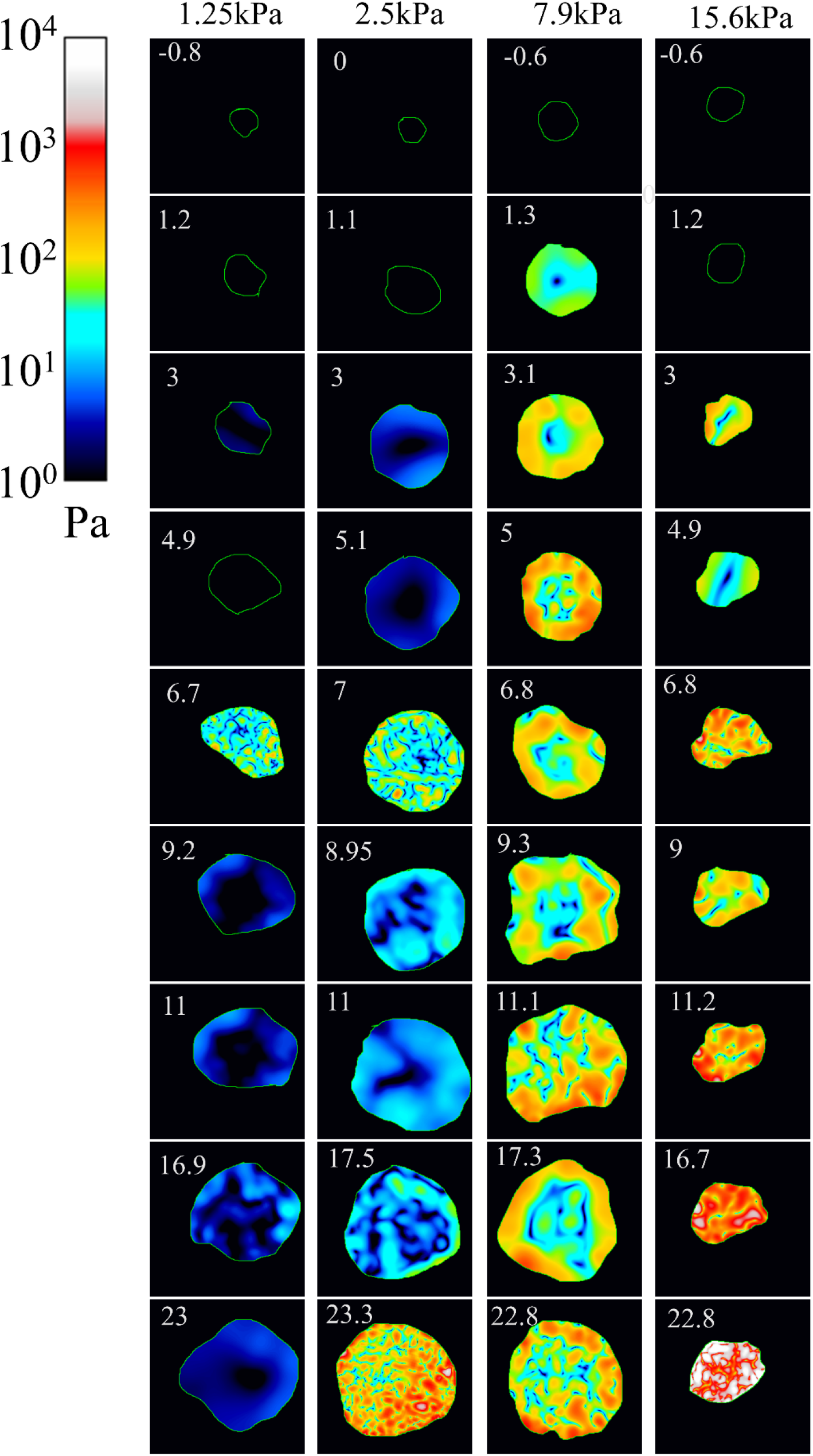
The phagocytic cup is mechanically organized: the mechanical stress is generated in delimited stress zones accumulating into a peripheral force ring. The mechanical stress of the four representative cells from Figure 4 is shown at selected time points as heat maps on a log color scale. The time points span the entire course of frustrated phagocytosis with 0 min being the beginning of spreading. The yellow outline represents the boundaries of the cell. Note that the color scale is the same for all conditions. These cells correspond to Movies 1 to 4.

**Figure 6:**
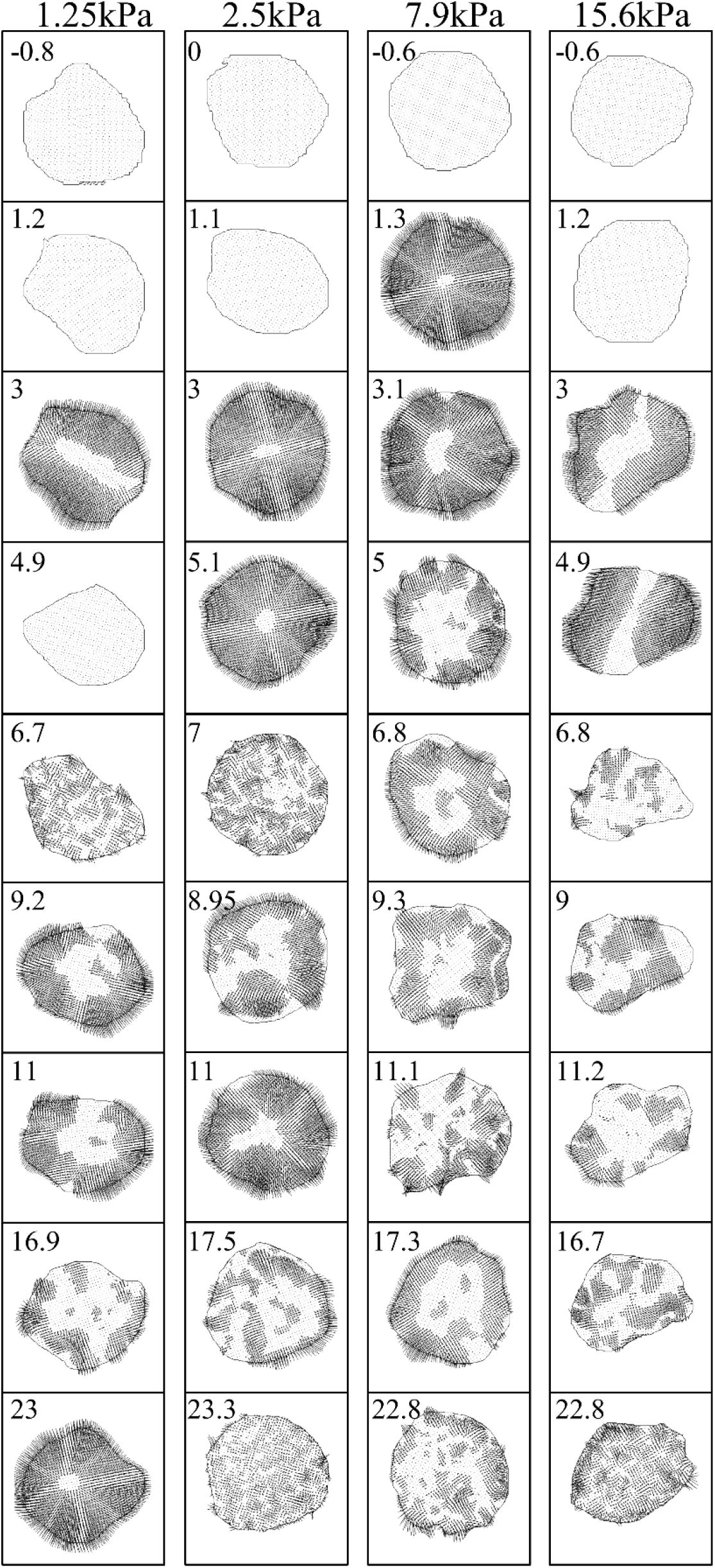
the forces generated during frustrated phagocytosis are centripetal. Mechanical stress vector maps corresponding to the stress magnitude maps shown in figure 5. The direction of the arrows indicates the orientation of the force exerted by the cell onto the substrate at this point.

**Figure 7:**
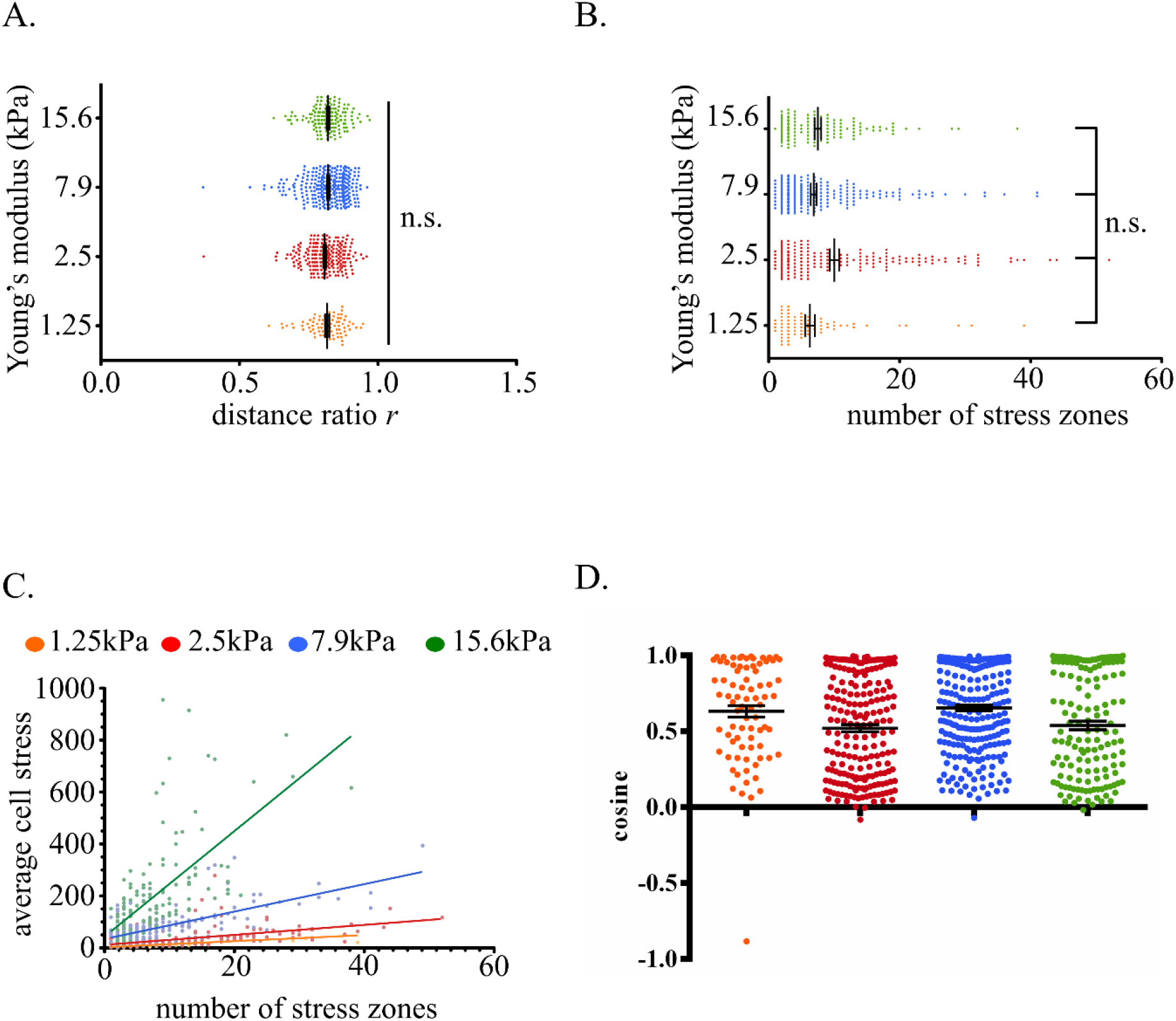
the characteristics of stress-generating zones are comparable between the substrate stiffness conditions. (A) For each time frame of each cell, the radial positions of all the detected stress zones (defined as the top 20% pixels on the stress map) were expressed as radial index *r* and averaged together. The index *r* approximates 1 for peripheral positions and 0 in case of a central position. (B) The number of separated stress zones, indicating the level of fragmentation of the stress map is comparable between the four stiffness condition. Each dot represents the number of stress zones of one given cell at one given time point. (C) The level of fragmentation in the stress map is positively correlated with the average amount of stress generated in that cell overall. The effect of stress map fragmentation on the average stress magnitude is more pronounced as the gel gets stiffer. (D) The centripetal orientation of the traction forces was assessed by the cosine of the angle between the traction force vector at a given point of the cell and the vector from that point to the centroid of the cell. As such, a cosine of 1 indicates a perfectly centripetal force, a cosine of −1 a perfectly centrifuge force and a cosine of 0 a perpendicular force. Cosines of all the force vectors of a cell at a given time point were averaged together which was then averaged for all the time points for each cell and shown as individual dots. The average ±SEM of all cells for each stress condition are shown in black. Orange = 1.25, red = 2.5, blue = 7.9 and green = 15.6 kPA as in the previous figures. Note that an averaged value of 0 correspond to a random orientation of the traction vectors.

### Mechanosensing and the subsequent modulation of stress generation are independent from tyrosine phosphorylation and actin polymerization

Our quantification of the cell mechanical stress showed that macrophages were able to respond to substrate stiffness. Tyrosine phosphorylation and actin polymerization downstream of FcγR ligation are both necessary for phagocytic cup formation (Garcia-Garcia & Rosales 2002). Cellular mechanical adaptation to increasing stiffness could thus be mediated by an increase in signal intensity resulting from greater tyrosine phosphorylation and/or higher F-actin concentration. To determine if this occurred we monitored global tyrosine phosphorylation (pY) and F-actin content in BMMs 10min after the onset of frustrated phagocytosis on either soft (2.5kPa PA gel) or stiff (15.6kPa PA gel or glass) substrates, a time at which BMMs have reached the stable phase of mechanical stress at this time (fig 1). In spite of a more than 2-fold increase in pY over that time, the pY level and F-actin content were not statistically significantly different between the stiffness conditions at either the onset (fig 8A) or after 10 min (fig 8B). This suggested that the sensing of target stiffness and its translation into a modulation of force generation was independent from the amount of phosphotyrosine-based signaling or F-actin generation.

**Figure 8:**
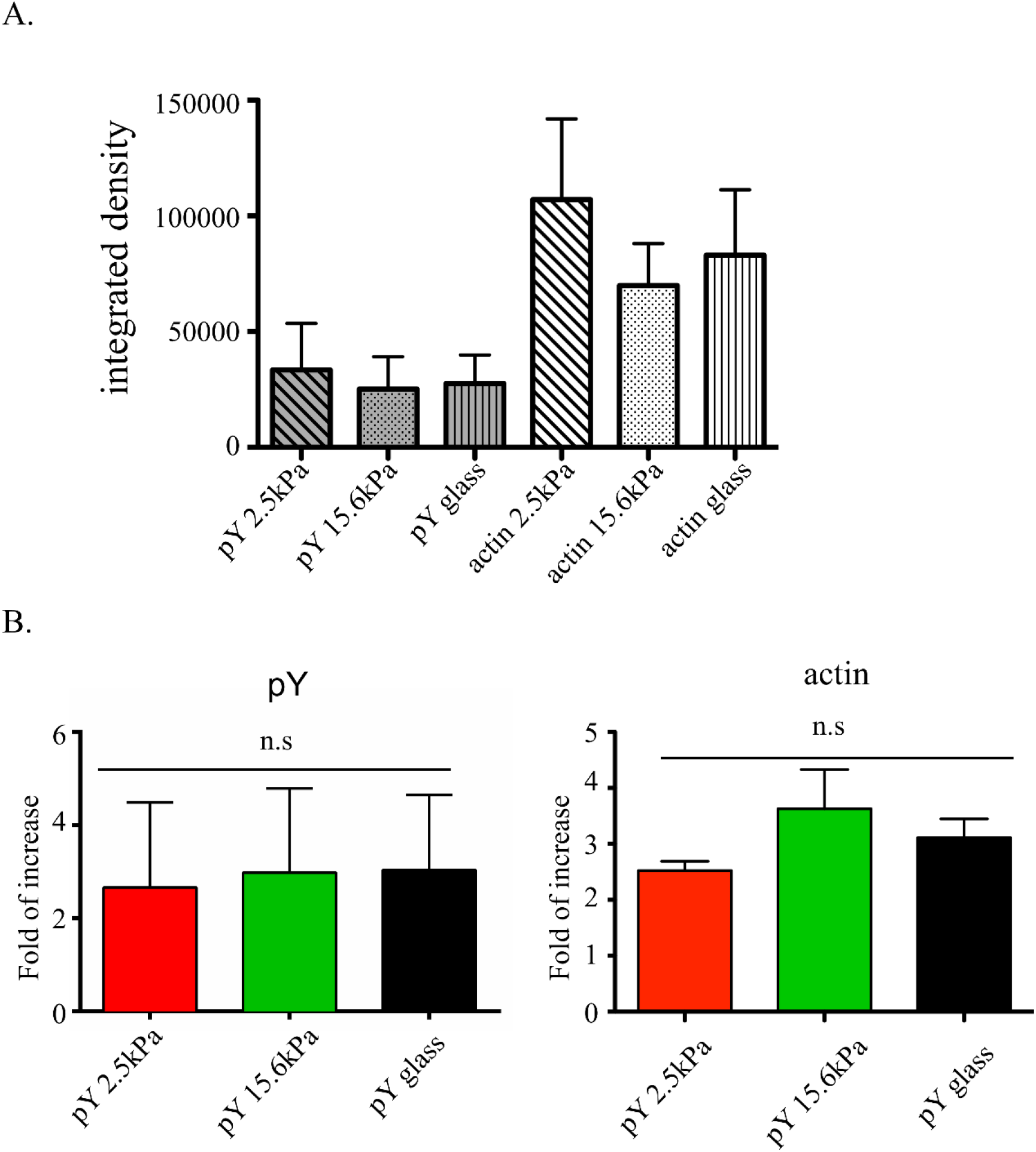
The mechano-adaptation of BMMs to target stiffness during frustrated phagocytosis is not dependent on pY or F-actin production. Quantification by immunocytochemistry of the total amount of pY and F-actin in BMMs. (A) The amounts are calculated by quantifying the cells integrated density. (A) pY and F-actin levels before initiation and (B) after 10min of synchronized frustrated phagocytosis on glass, 2.5kPa gel, or 15.6kPa gel. Only the cells showing a clear initiation of frustrated phagocytosis (cell area reaching at least 600um^2^) are considered and normalized to the average integrated density of pY or F-actin at t = 0min.

## DISCUSSION

Phagocytosis is a widespread and multi-purpose process owing to the ability of macrophage to engulf targets of various sizes with different physical and chemical properties. We wanted to know how primary macrophages would respond to such a diversity of particle stiffness. We hypothesized that the mechanics controlling phagocytic cup formation was modulated as a function of the target stiffness. We modelled FcγR-dependent phagocytic cup formation by studying macrophage spreading on an IgG-coated substrate. Frustrated phagocytosis is a useful and commonly used proxy of “real” phagocytosis. Indeed, even though this approach does not recapitulate important aspects of cup formation such as its 3D shape, the cell basal membrane shares strong molecular similarities with phagocytic cups and is compatible with TFM.

We quantified the kinetics of cell deformation as well as the spatiotemporal dynamics and magnitude of cell-generated mechanical stress transmitted to the substrate during frustrated phagocytosis. Our principal finding is that the magnitude of mechanical stress generated downstream of FcγR engagement is linearly proportional to the substrate while the associated spreading response remains strikingly unchanged. This conservation of the membrane extension rate over different substrate stiffness is clearly different from what has been reported in the case of non-phagocytic cues such as collagen (Engler et al 2004, Li et al 2014, Yeung et al 2005), fibronectin (Han et al 2012, Nisenholz et al 2014, Tee et al 2011, Vishavkarma et al 2014), laminin (Reimer et al 2017) where spreading is correlated to substrate stiffness. Our interpretation is that during frustrated phagocytosis, macrophages are able to conserve the same deformation velocity and efficiency thanks to a mechano-adaptation to the target rigidity. This mechano-adaptation is characterized by a modulation of the force magnitude and a preservation of the time kinetics and spatial organization.

Our force maps show that the stress generation is fragmented into distinct stress zones. The stress zones accumulate at the cell periphery, suggesting that the signaling and cytoskeleton proteins responsible for mechanical force generation were also localized to the outer edge. Consistent with this, the phosphoinositide PI(4,5)P_2_, the activity of the Rho GTPase Cdc42 and the downstream polymerization of actin have been shown to be spatially restricted to rims of the cup (Hoppe & Swanson 2004, Swanson 2008). Similarly, it has recently been shown that the rim of actin is made of short-lived podosome-like structures, that disappear from the basal/central area upon PI(4,5)P2 hydrolysis (Ostrowski 2019). Our spreading kinetics during frustrated phagocytosis appeared similar to what has been reported for a monocyte/macrophage cell line (Masters et al 2013) and for fibroblasts or endothelial cells (Dubin-Thaler et al 2008, Reinhart-King et al 2005). A first phase of rapid linear spreading, P1, followed by a second phase of slowing non-linear spreading, P2. The spreading curves stop fitting a linear regression after 7min consistent with observations in other models of spreading with a possible over-estimation due to the time resolution used. It has recently been reported that the J774.1 cell line exhibited a marked cell retraction at the end of frustrated phagocytosis associated with a transient peak of traction forces (Kovari et al 2016). However, we observed neither a conspicuous transient peak of force at the end of spreading nor a subsequent cell retraction. Such strong retraction has not been reported in other studies from other groups either (Beningo & Wang 2002, Cannon & Swanson 1992, Masters et al 2013). These behavioral differences may be due to the difference of cellular model; in our case, we used primary cells rather than cell lines.

We noted that spreading during frustrated phagocytosis begins before any detection of mechanical stress. This may be explained by the presence of low level stresses that are below the sensitivity of our assay. Alternatively, it can be interpreted as an initial “jump out” of the membrane. Although contact with the substrate is needed to trigger signaling downstream of the FcγR, the membrane would then extend isotropically without strong anchoring to the substrate, in contradiction to the classic “zipper model” (Swanson & Baer 1995). Further extension on the contrary may require firm binding to the substrate and increase in force. This explains at the mechanical level the observation made by Master et al. that macrophages can extend protrusions of non-opsonized regions at early time points. At later time points however, anchoring becomes necessary and protrusions become restricted to opsonized areas (Masters et al 2013). Consistent with our results, the initial phase of neutrophil spreading is almost certainly due to the release of the cortical tension, since they found no effect of myosin II inhibitors on the outward propagation of cell spreading (Henry et al 2015). We also observed that although strongly variable, stress rapidly stabilized around a non-null value while the cell remains in a mechanical non-equilibrium and keeps spreading. Such a time lag between stress and strain is reminiscent of the viscous creep of viscoelastic systems (Kollmannsberger & Fabry 2009, Kollmannsberger & Fabry 2011) and suggests a key role of rheology in regulating frustrated phagocytosis. Finally, we did not observe a correlation between the magnitude of stress generated and cell spreading velocity. Altogether our results suggest that the generation of mechanical stress is necessary to force the deformation of the cell membrane into a phagocytic cup but does not control the kinetics and the extent of the cup growth.

The mechanoresponse that we report herein is in agreement with other works demonstrating a correlation between cellular mechanics and environmental physical properties, where substrate stiffening correlated with increased cell stiffness or force generation (Crow et al 2012, Hind et al 2015, Jannat et al 2011, Labernadie et al 2014, Paszek et al 2005, Webster et al 2014). However, the mechanics of phagocytic cup formation holds some unique features. First, macrophage FcγR phagocytic response is stronger than that observed during neutrophils phagocytosis (up to 600nN for the former, 30nN for the latter (Herant et al 2005)) but weaker than macrophage migration on fibronectin (up to 3000nN in that case (Hind et al 2015)). Second, the mechanical stress appears to stabilize *before* the cell reaches its spreading plateau. By contrast, fibroblasts only generate very low mechanical stress during spreading and only display a marked and sudden increase once they have reached their maximal area (Dubin-Thaler et al 2008). Third, phagocytosis occurs at a much faster rate. While force measurements of macrophage migration occur over several hours (Hind et al 2015) it only takes a few minutes for the cells in frustrated phagocytosis to sense substrate stiffness, deliver a spatially structured mechanical stress and reach several folds of increase of their area.

The existence of rapid macrophage mechanosensing had been reported in an previous study, albeit without time-resolved quantification of cell deformation and mechanical stress (Beningo & Wang 2002). The authors reported that macrophages were more efficient at performing frustrated phagocytosis on harder PA substrates. Our own conclusion differs in that the stiffness of the target has an impact on the cell mechanics only while the morphological dynamics remains invariant. In contrast to our PA gels functionalized by direct covalent linking with IgG, Beningo *et al.* used a combination of crosslinked BSA and anti-BSA rabbit IgG. We observed less engagement and spreading on substrates coated with BSA and IgG (data not shown). It is therefore possible that our covalently linked IgG-only coating results in an overall stronger signal and allows similar phagocytosis efficiency on different substrate stiffness.

Our work raises two fundamental questions. 1. What are the molecular mechanisms generating the additional force on stiffer substrates? Our pY and F-actin staining suggested that increased signaling of and bulk actin polymerization may not be an explanation for the increased force. Another possible explanation is that auxiliary molecular motors are mobilized on stiffer substrates. Many different myosin family members have been detected at the phagocytic cup including myosins II, X, V, IXb, Ig and Ic (Araki et al 2003, Cox et al 2002, Dart et al 2012, Swanson et al 1999). However, their precise mechanical contribution to phagocytic cup formation remain unclear and further work is needed to determine their potential involvement in tuning of stress magnitude constitutes an important next step for investigation. 2. How do macrophages sense the phagocytic target stiffness? Mechanosensing necessitates an association between the cytoskeleton and a mechanoreceptor so as to probe the substrate, as it is the case with focal adhesions (Humphrey et al 2014). A possible candidate could be the Myosin I family. Indeed, Myosin IE,F,G all connect the membrane to the cytoskeleton, are present at the phagocytic cup and have been shown to participate in target engulfment (Barger et al 2019, Dart et al 2012). Alternatively, the presence of adhesive integrins and podosome-like structures may also contribute to mechano-signaling (Maxson et al 2018, Ostrowski et al 2019). While the FcγR is classically seen as a signaling receptor, comparative reasoning suggests that it might also act as a mechanoreceptor. The FcγR is an ITAM-based receptor similarly to the T Cell Receptor (TCR), that has been reported to trigger force generation in T lymphocytes and to mediate mechanosensing of the stiffness of antigen presenting cells (Judokusumo et al 2012, O’Connor et al 2012). The TCR works as a bridge between the cytoskeleton and the outside through its association with actomyosin (Comrie & Burkhardt 2016). It is thus possible that the FcγR works in the same way. Whether the FcγR is able to connect with the actin network, probe the mechanical features of the phagocytic target stiffness and to feed that information back into the cell is an important question worthy of further investigation.

## CONCLUSION

We have shown that primary macrophages demonstrate a mechano-adaptation to their target during FcγR-triggered frustrated phagocytosis: the amount of mechanical stress generated is linearly correlated with the stiffness of the target. These mechanical sensibility and adaptability are associated with a striking invariance of the cell deformation between different target stiffness. It may thus constitute the mechanical basis that explains the ability of macrophage to face physically diverse targets. We also observed that frustrated phagocytosis seems to be initiated without significant traction onto the substrate before requiring anchoring to sustain membrane extension. This mechanical analysis of macrophage phagocytosis thus brings us new physical measurements useful for *in silico* modeling as well as novel and valuable information to our understanding of phagocytosis. Identifying the molecular sensors and motors responsible for the assessment of stiffness and force modulation would be a key development to harnessing phagocytosis for biotherapeutic use.

## ACKNOWLEGEMENTS

We thank the members of the Condeelis, Segall and Hodgson laboratories for helpful comments and discusions. We thank Samer Hanna and Veronika Miskolci for their assistance with the experiments. We thank Daniel Hammer and Laurel Hind for their help and guidance in the TFM experiments. We thank Micah Dumbo for having authorized the use of his LIBTRC software. We thank Kessler McCoy-Simandle, Nathan Bessa Viana, Karine Anselme and Allison Sharon Harney for editorial assistance with the manuscript. This work was supported by the NIH grant GM071828 to Dianne Cox.

## SUPPLEMENTARY FIGURES AND TABLES

**Supplementary Figure S1:**
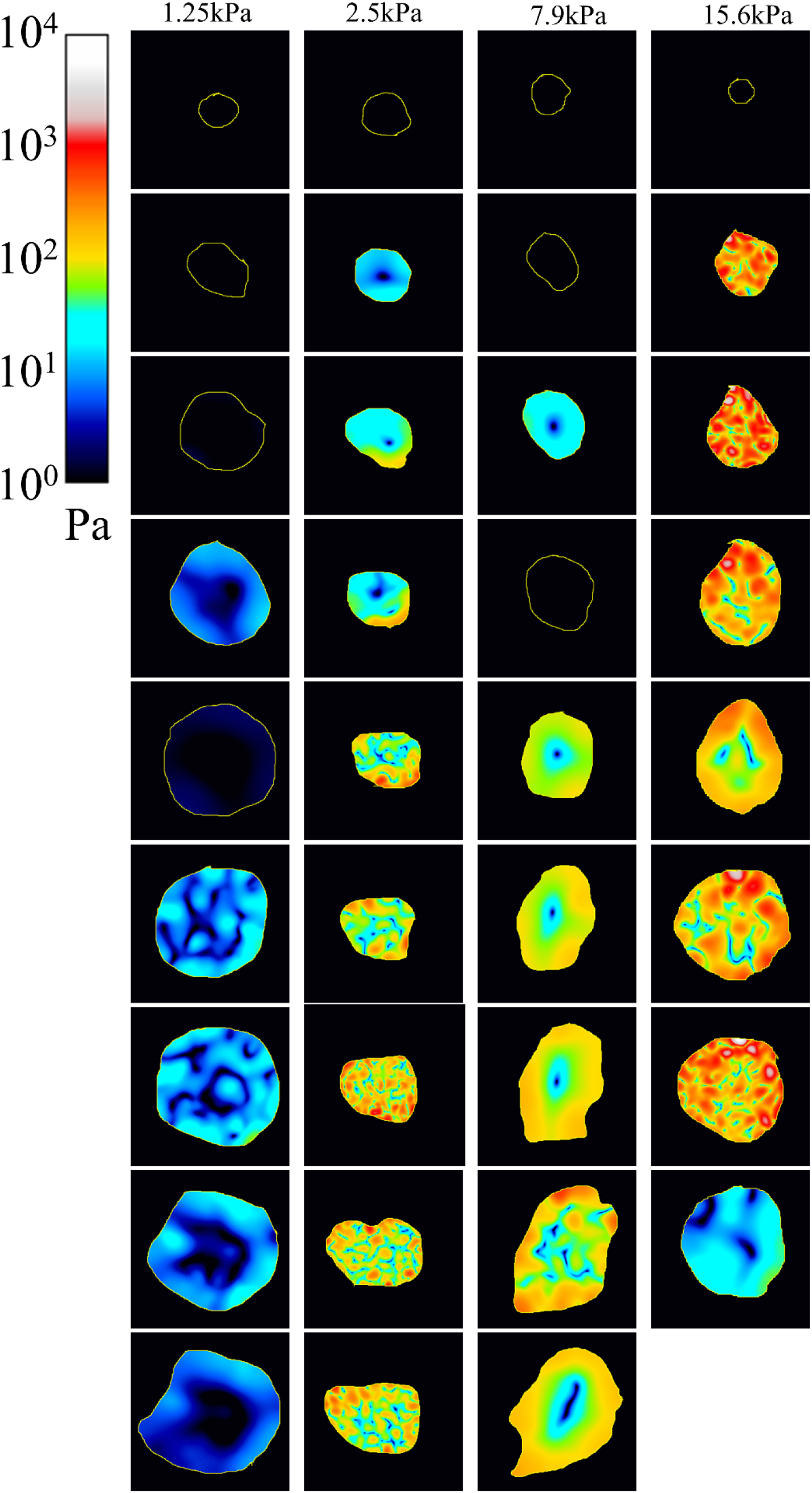
additional examples of stress map during frustrated phagocytosis. The mechanical stress of 4 additional representative cells (one per stiffness condition). Note that the log color scale is the same. Time is indicated in minutes. These cells correspond to Movies 5 to 8.

**Supplementary Figure S2:**
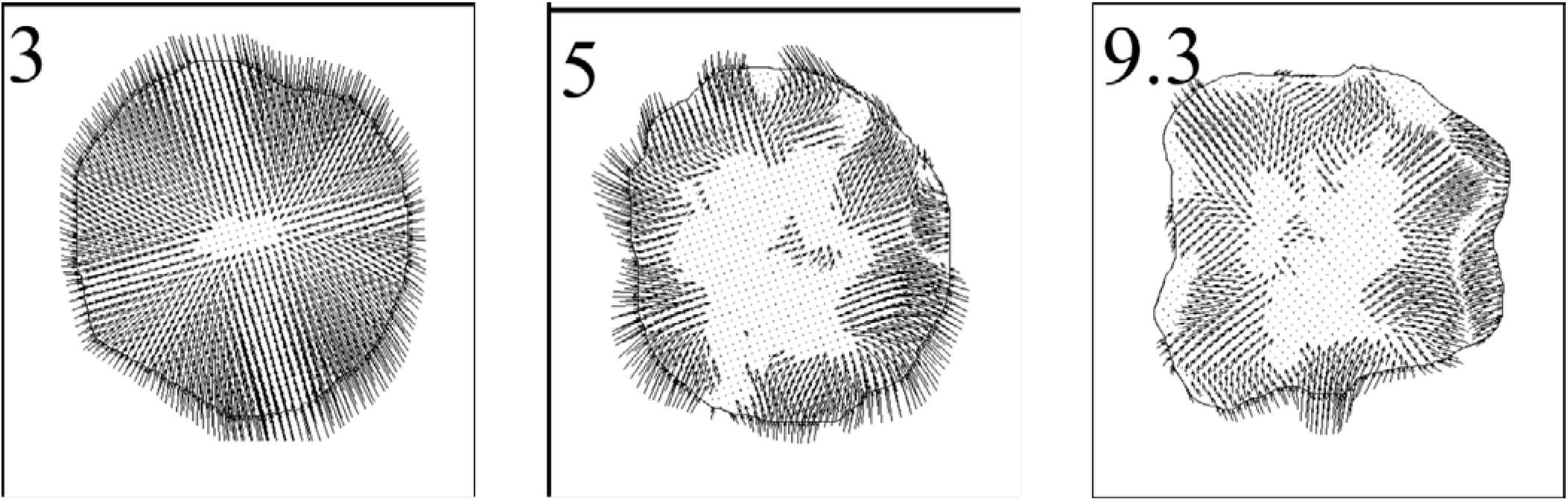
close up of stress vector map from Figure 6. Close up of 3 force vector map from Figure 6 highlighting the centripetal organization of force vectors

### MOVIES

For all movies, the time (minutes) is set with t = 0min at the onset of frustrated phagocytosis.

**Movie 1-4**: Full time course of frustrated phagocytosis for the representative cell spreading over a 1.25kPa, 2.5kPa, 7.9kPa, 15.6kPa (respectively) gel shown in Figure 4,5,6. Left: phase image, Right: stress map. Cell outlines are shown in yellow.

**Movie 5-8**: Full time course of frustrated phagocytosis for the representative cell spreading over a 1.25kPa, 2.5kPa, 7.9kPa, 15.6kPa (respectively) gel shown in Supplementary Figure S1. Left: phase image, Right: stress map. Cell outlines are shown in yellow.

